# Magnetic actuation of otoliths allows behavioral and brain-wide neuronal exploration of vestibulo-motor processing in larval zebrafish

**DOI:** 10.1101/2023.04.18.537408

**Authors:** Natalia Beiza-Canelo, Hippolyte Moulle, Thomas Pujol, Thomas Panier, Geoffrey Migault, Guillaume Le Goc, Pierre Tapie, Nicolas Desprat, Hans Straka, Georges Debrégeas, Volker Bormuth

## Abstract

The vestibular system in the inner ear plays a central role in sensorimotor control by informing the brain about the orientation and acceleration of the head. However, most experiments in neurophysiology are performed using head-fixed configurations, depriving animals of vestibular inputs. To overcome this limitation, we decorated the utricular otolith of the vestibular system in larval zebrafish with paramagnetic nanoparticles. This procedure effectively endowed the animal with magneto-sensitive capacities: applied magnetic field gradients induced forces on the otoliths resulting in robust behavioral responses comparable to that evoked by rotating the animal by up to 25°. We recorded the whole-brain neuronal response to this fictive motion stimulation using light-sheet functional imaging. Experiments performed in unilaterally injected fish revealed the activation of a commissural inhibition between the brain hemispheres. This magnetic-based stimulation technique for larval zebrafish opens new perspectives to functionally dissect the neural circuits underlying vestibular processing and to develop multisensory virtual environments, including vestibular feedback.

**Highlights:**

- After injecting a ferrofluid into the inner ear of a larval zebrafish, the ear-stones can be actuated via magnetic forces.
- This method allows one to elicit vestibular-like behavioral responses without impairing physiological inner ear functions.
- It is compatible with brain-scale functional imaging and thus offers a promising avenue to investigate the neural underpinnings of vestibular-driven behaviors.

**eTOC:** Ferrofluid injection into zebrafish inner ear allows magnetic manipulation of ear-stones to evoke vestibular responses in static animals. This in vivo method is compatible with brain-scale imaging, offering a promising approach to investigate neural mechanisms underlying vestibular-driven behaviors.

## 1 INTRODUCTION

The vestibular sense is crucial for evidence accumulation and decision-making in every ethological behavior that involves body movements. In vertebrates, the vestibular apparatus is located in the inner ear and comprises several organs that sense head/body movements and sound. In mammals, two otolith organs: the saccule and the utricle, together with the semicircular canals, transduce linear and angular acceleration, whereas the cochlea transduces auditory stimuli. Fish do not have a cochlea; instead, the saccule and likely the lagena have evolved to transduce water vibrations in the auditory frequency range^1,2^. Rotational acceleration of the head induces an endolymph flow in the semicircular canals, which is detected by mechanosensitive structures called cupulae. Translational acceleration, as well as gravitational forces, act on the otolithic structure overlaying on the utricular and saccular epithelia, and whose motion is transduced by mechanosensitive hair cells to which they are coupled.

Neuronal signals encoding the head orientation and movement are relayed to neuronal circuits that drive compensatory movements in order to stabilize gaze and posture. Vestibular information is first processed in the brainstem vestibular nucleus and the cerebellum, which receive direct vestibular afferent input. Information is further distributed to oculomotor, skeletomotor, and autonomous motor systems, and in mammals, also via the thalamus to cortical systems^3^. At the various stages of signal processing, vestibular information is integrated with non-vestibular signals of self-motion information such as visual, somatosensory and proprioceptive inputs as well as with locomotor efference copies^4^.

In spite of the central role played by the vestibular system in sensorimotor tasks, most neuronal recordings are currently performed in animals deprived of any vestibular signals, i.e., under head- or body-fixed stationary conditions. This is due to the inherent challenge of combining neural recordings and natural vestibular stimulation as the latter necessitates to rotate or translate the animal’s head in space, and is thus incompatible with head-fixed recording configurations required for most functional calcium imaging techniques or electrode recordings. Our knowledge of the vestibular system thus essentially derives from electrophysiological experiments in which the spike activity of a few neurons is sequentially monitored using implanted electrodes.

As vestibular processing is widely distributed across the brain, zebrafish constitutes a promising model animal to study the neuronal substrate of this highly conserved sensory system. The small size and transparency of the larval zebrafish brain indeed offers the unique opportunity to record cell-resolved brain-wide neuronal activity using light-sheet based calcium imaging^5,6,7,8^.

As early as 6 days post-fertilization^9^, an age at which whole-brain imaging is routinely performed, larvae efficiently stabilize their posture and gaze in response to body rotation, via vestibulo-ocular and vestibulo-spinal reflexes^9,6,10,11,12^. The semicircular canals in larval zebrafish are not yet functional at 6 dpf,^13^ so all head motion signals are solely transduced by the utricle at this age. This makes the interpretation of vestibular stimulation easier in larval zebrafish compared to later developmental stages or to other vertebrate species, where otolith organs and semicircular canals contribute simultaneously to the transduction of head motion into nerve signals.

Experimental methods to provide controlled vestibular stimulation, while performing functional calcium imaging in larval zebrafish brain, were recently introduced. In the work by Migault *et al.*^6^, we solved the problem by co-rotating the fish and the (miniaturized) light-sheet microscope, thus keeping the imaging volume unchanged while providing a physiological vestibular stimulation. Favre-Bulle et al.^10,7^ generated a fictive vestibular stimulus using optical tweezers to displace the utricular otolith, with the advantage that it can target a single otolith. Hamling et al.^14^ tilted quickly the fish in a platform and imaged right after the platform came back to the horizontal position. These methods are very powerful. The first two enable simultaneous neural recording, although both involve demanding optical developments that may hamper their broad diffusion among groups employing neurophysiological methods. Furthermore, the accessible stimulation range, in terms of maximal acceleration that they can emulate, is limited. The last method does not allow recording neuronal activity during the time course of a dynamic vestibular stimulus.

Here, we present an alternative approach based on the magnetic actuation of the otoliths after surface coating by ferromagnetic nanoparticles. These superparamagnetic iron oxide nanoparticles are available in the form of colloidal solutions called ferrofluids^15^. Although lacking a permanent magnetic moment, these particles acquire a magnetization in an externally applied magnetic field and can be manipulated by magnetic field gradients. Their magnetic susceptibility is several orders of magnitude larger than that of biological tissues^16^, allowing the application of large forces. Biocompatible ferrofluids have been used to study mechanical properties inside living tissues *in vivo*^17,18,19,20,21^ and functionalized nanoparticles have allowed targeting cellular components such as DNA and proteins with high specificity^22,23^ or to deliver drugs into compartments that are difficult to access as, e.g., the inner ear^16^.

Magnetic actuation offers several advantages over optical tweezers in the context of biological systems. First, biological tissues are fully transparent to magnetic fields. Forces can thus be exerted in a controlled way deep within the specimen, regardless of its optical transparency. Second, magnetic fields do not induce heating, and, except for magnetoreceptive species^24^, most animals are insensitive to this physical parameter. Thus, besides the injection itself, this technique is physiologically non-invasive even for extremely large magnetic intensities.

Here, we show that the injection of ferrofluid into the otic vesicle of larval zebrafish allows controlled magnetic forces to be exerted on the otolith, mimicking naturally occurring gravitational and acceleration forces. This fictive vestibular stimulation elicits strong and robust compensatory eye and tail movements, comparable to those evoked by roll or pitch tilts of the animal over large angles. We simultaneously recorded the brain-wide neuronal activity evoked by this fictive vestibular stimulation using functional light-sheet microscopy. By injecting the ferrofluid into a single ear, we disentangled the contribution of each utricle to the brain-wide neuronal response, which is not possible under natural conditions when rotating the animal^6^ during whole-brain imaging but has been successfully performed using optical tweezers [10, 7]. This constitutes the first use of a ferrofluid to stimulate a sensory system *in vivo*. The method is inexpensive, easy to implement and compatible with most neurophysiological recording methods such as large scale functional imaging, optogenetics or electrophysiology. This makes vestibular stimulation now broadly accessible and will allow researchers with different research foci to investigate the vestibular contribution to a large variety of behaviors and neuronal processes such as evidence accumulation, head direction circuits, decision-making, cerebellar functions, motor learning and motor control and this in head attached or eventual even freely swimming assays.

## 2 RESULTS

### After ferrofluid injection into the otic vesicle, vestibular-driven behaviors can be evoked through magnetic stimulation

We injected a custom-made ferrofluid^25^ into both inner ears of zebrafish larvae 5 days after fertilization (dpf). The ferrofluid consisted of 11 nm in diameter iron oxide (γ-Fe_2_O_3_) particles with citric acid surface functionalization to make them stable in water due to their negative surface charge (pH 7, see Methods). Similar ferrofluids are also commercially available (see the discussion section and the Star Methods). After the injection, the otic vesicle maintained its shape and the ferrofluid was visible as a red-orange tinge (Figure 1A). The utricular otolith itself, once dissected out and washed, also displayed a slight orange coloration, indicating that some injected ferrofluid particles had permanently bound to the otolith. The otolith thus became magnetically actuable, as confirmed by approaching a permanent magnet in its proximity. The otolith immediately moved towards the magnet, as shown in the Supplementary Figure S1A. The same is likely true for the otolith of the saccule. However, in this study, we focused only on the utricle.

**Figure 1:**
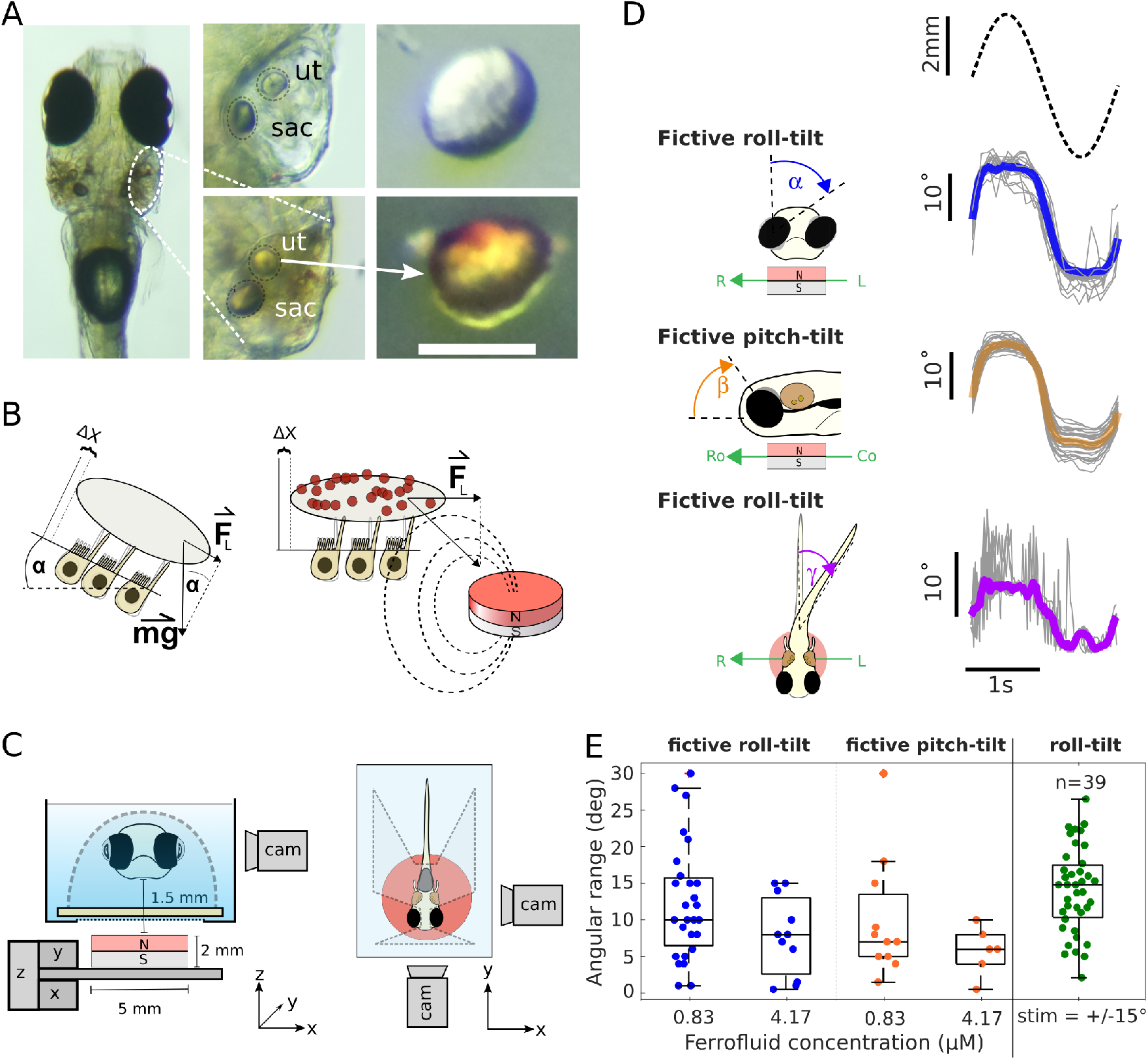
Magnetic actuation of the otoliths after surface coating by ferromagnetic nanoparticles. **A.** Top view of a 5 dpf zebrafish larva after bilateral intra-otic ferrofluid injection. Middle column: Zoom onto the otic vesicle with injected ferrofluid (bottom) and before injection (top). The otoliths of the utricle, ut, and of the saccule, sac, are visible. Right: Bright-field image of a utricular otolith dissected from a control fish (top) and from a fish after ferrofluid injection (bottom). Attached iron nanoparticles appear in red-orange. Scale bar = 50 µm. **B** The diagram on the left illustrates the lateral force experienced by an otolith when the head is rotated relative to the gravitational field, F*_L_* = g *·* sin(α). The right side illustrates an otolith covered with nanoparticles that exert a lateral force onto the otolith when placed in a magnetic field gradient. **C** Left: Diagram of the setup in front view. An x,y-motorized stage and a manual z-stage (gray) move the magnet (red) under the head-tethered fish, mounted in agarose (outlined by a dashed line). Right: Top view of the setup, illustrating the front and side cameras (cam) for eye motion tracking. The magnet center is aligned with the center of mass of the fish inner ears (not drawn to scale). Caption continues next page. **D** Eye and tail behavioral responses elicited by a magnet that moved in different directions beneath a fish. Fish were injected with 0.83 µM ferrofluid solution. The magnet moved sinusoidally at 0.5 Hz and 2.5 mm in amplitude for one minute corresponding to 30 cycles. Shown are one period of the stimulus (dashed line), the mean responses (color) and the responses per cycle (gray). Cycles where tracking failed were removed, resulting in N = 27, 30, 29 respectively. Note that we chose the stimulation amplitude of 2.5 mm to exceed the linear range of the force-displacement relationship (see Figure 3) in order to apply the maximum possible lateral force to the otolith. This explains the apparent saturation of the behavioral responses. The variability in the tail response reflects transiently evoked swim bouts. **E** Angular range of evoked eye rotation angles (peak-to-peak) in response to fictive roll and pitch-tilt stimuli plotted for two concentrations of ferrofluid bilaterally injected into the inner ears (N*_roll,_*_0.83_ *_µM_* =27, N*_roll,_*_4.17_ *_µM_* =11, N*_pitch,_*_0.83_ *_µM_* = 11 and N*_pitch,_*_4.17_ *_µM_* =6). The measured behavioral responses at the two concentrations are neither significantly different for the roll nor for the pitch axis (p*_roll_* = 0.11, p*_pitch_* = 0.36, Kruska-Wallis test). The last box plot shows the eye motion of 39 fish to a physiological sinusoidal roll stimulus of *±*15° in amplitude. See also also Figure S1, Video S1,, Video S2, and Video S6.

Next, we tested whether this nanoparticle coating of the otolith could yield magnetic forces *in vivo* on the otolith comparable to the gravitational force that acts on it when the head/body is roll- or pitch-tilted in space (Figure 1B). To do so, we examined the behavioral response (compensatory eye and tail movements) that were induced through magnetic actuation. We thus immobilized a bilaterally injected fish in a drop of 2% low melting point agarose on a thin glass slide and removed the agarose around the eyes and tail to allow free movements. By hand we imposed in-plane movements of a small permanent neodymium magnet beneath the fish. When the magnet was moved along the medio-lateral axis, the eyes rolled and the tail bent in a direction opposite to the magnet and discrete swim bouts were evoked (Video S1, Part I). Such movements are characteristic of responses elicited by a roll motion of the animal (i.e., a rotation along its longitudinal axis) via vestibulo-ocular and vestibulo-spinal reflexes^6,9^. In this case, the magnetic force acted laterally on the otolith, as does the gravitational force during a roll motion. When the magnet was moved along the antero-posterior axis, the eyes rotated along the pitch axis and discrete swim bouts were triggered. Here the response to the magnetic stimulation was in line with compensatory eye movements and tail kinematics elicited upon pitch-tilting the fish^9^ (see Video S1 Part II).

To perform quantitative experiments, we automatized the system, installed front and a side cameras to better monitor the behavioral responses and mounted this stimulation unit on a light-sheet microscope. We imposed controlled sinusoidal displacements to the magnet using a two-axis motorized stage, either along the lateral axis (fictive roll-tilt stimulus) or along the antero-posterior axis (fictive pitch-tilt stimulus). We used a frequency of 0.5 Hz and an amplitude of 2.5 mm corresponding to the radius of the magnet to reach the maximum of the magnetic force (see the following section describing the numerical simulations for an estimation of the corresponding magnetic force). We quantified the responses for a concentrations of 0.83 µM and 4.7 µM of injected ferrofluid. Typical behavioral responses recorded at 6 dpf (one day after the injection of the low concentrated ferrofluid) are shown in Video S2 (Part I-III) with minimal cross-talk between the two stimuli directions, provided that the magnet was well centered beneath the fish. The method was robust: out of 27 successfully injected fish, 23 responded with a peak-to-peak eye motion angle larger than 5 degrees to a moving magnet in the roll axis (Figure 1E). The observed vestibular behaviors were reproducible and stable over time, with only small variability over 30 stimulus repetitions in the same fish (Figure 1D). From the averaged cyclic ocular rotation signal, we extracted an angular range of α = 12.4° *±* 7.8° (mean *±* standard deviation, N=27) during simulated roll-tilt and β = 10.3° *±* 8.5° (mean *±* standard deviation, N=11) for simulated pitch-tilt (Figure 1). Using a higher ferrofluid concentrations did not result in significantly different responses(Figure 1E). In the following, we used the lower concentrated ferrofluid solution for all experiments. Importantly, stimulating the fish with the magnet at 9 dpf, four days after the injection, still evoked the same behavioral responses (see Figure S1 C). This demonstrates that the injected ferrofluid remained stable in the ear for several days post-injection. And similar behavioral responses were successfully evoked also with a commercially available ferrorfluid (see Figure S1C).

These values can be compared to those obtained during natural vestibular stimulation, in which the animal is actually roll- or pitch-tilted in space. As an illustration, we show in Figure 1E the angular range (α = 14.1° *±* 5.8° (mean *±* standard deviation, N=39) of the eye rotation measured in larvae exposed to a sinusoidal roll motion of *±*15°. The variance of the data in both experiments are comparable (p=0.09, two-sample F-test), indicating that the large variability across specimens is not specific to the fictive ferromagnetic stimulation. From this roll-tilt evoked responses under natural vestibular stimulation, we calculated a gain of the vestibulo-ocular reflex in darkness for the roll-till direction of g*_roll_* = 0.47 *±* 0.19 (mean *±* standard deviation, N=39). For the pitch-tilt direction a gain of g*_pitch_* = 0.3 was reported^9^. From this calibration, we thus estimated that the fictive magnetic vestibular roll and pitch-tilt stimuli corresponded to a peak-to-peak stimulus of α/g*_roll_ ≈ ±*26.38° *±* 4.88° (mean *±* SEM) and β/g*_pitch_ ≈ ±*29.42° *±*10.24° (mean *±* SEM), respectively. We confirmed this observation by submitting injected fish to sinusoidal physiological roll stimuli of 20 and 30° peak-to-peak amplitudes, and subsequently to a fictive roll-tilt stimulation. The fictive stimulus evoked eye movements were comparable in amplitude to the ones evoked by the 30° peak-to-peak physiological roll stimulus consistent with our previous estimation (see Figure S1D). Furthermore, in the absence of a vestibular stimulus we did not observe neither a static bias nor a smooth drift of the orientation of the eyes (Video S2, Part IV) that could be an indication of a malfunction of the vestibular system.

We also tested the response to static roll stimuli in the form of angular steps of *±*2.5 mm magnet movements with a dwell time of 5 s and a transition time of 0.5 ms. Stepwise eye rotations were elicited (see Figure S1B), demonstrating that fast vestibular stimuli can be applied with this method. As observed during natural stimulation, the eyes did not remain static at the maximal eccentric angle but relaxed back due to an adaptation process most likely located in the inner ear. However, the adaptation was incomplete and a tonic response component remained. This residual eccentricity is an indication that we applied a direct force to the otolith because a drag force would be proportional to the velocity of the stimulus and would therefore only result in a phasic response.

### Ferrofluid injection into the inner ear does not impair vestibular function

Hair cells in the vestibular system are sensitive to mechanical and chemical stress, which can lead to cell death, thus impairing sensory function^26^. We assessed possible damage induced by either the injection procedure or by the ferrofluid itself using a simple behavioral assay. Fish use their vestibular system to keep their dorsal side-up posture stable during swimming. Therefore, uncorrected rolling along the rostro-caudal body axis during a swim bout can be used as an indicator for vestibular dysfunction^27,28,29^ (Figure 2A). We quantified the outcome of this procedure by calculating a roll ratio, i.e., the proportion of roll events over a total of 5 swimming events after a mechanically evoked startle response^30^ (see Methods). The roll ratio was measured at 2, 24 and 48 hours after the injection had been performed at 5 dpf.

**Figure 2:**
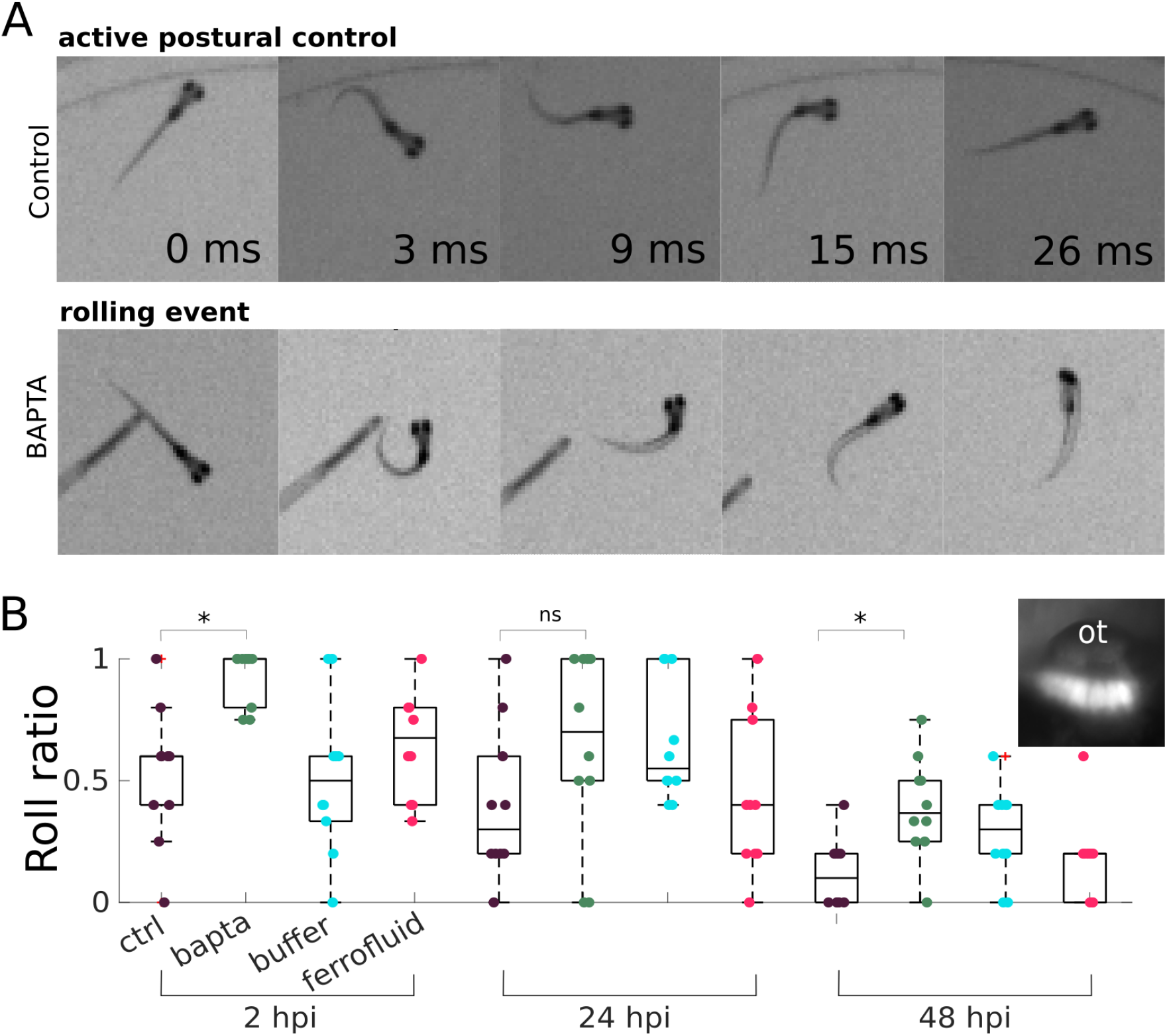
Control experiments probing the impact of the injection procedure on vestibular functionality. **A.** Image sequence recorded during evoked startle response behaviors for a non-injected control fish illustrating active postural control (top) as well as for BAPTA injected fish illustrating a roll event (bottom). **B** Roll ratio during an evoked startle response measured 2, 24 and 48 hours post bilateral injection (hpi) of BAPTA, ferrofluid or buffer compared to non-injected control fish from the same batch, respectively (N = 10 fish for each condition). The picture in the inset shows utricular hair cells labeled after the injection of 4-Di-2-ASP into the inner ear. See also Video S3, Figure S2, and Video S4.

Control (non-injected) fish had a mean roll ratio of 0.44 *±* 0.36 (mean *±* std, N = 10) at 5 dpf (Figure 2B). Although the vestibulo-ocular reflex is fully established at 5 dpf^9^, vestibular-driven postural control is still being refined between 5 to 7 dpf as evidenced by the decrease of the roll ratio during this period. As a positive control, we performed a similar assay on larvae injected bilaterally with the calcium chelator BAPTA (5 mM), which disassociates hair cell tip-links and disrupts the mechano-electrical transduction^31^. Two hours after the injection, the roll ratio was close to one (mean roll ratio: 0.93 *±* 0.11, see Video S3) and significantly different from the control (p=0.0078, Wilcoxon signed rank test), indicative of an almost complete loss of vestibular-driven postural control. At 48 h after BAPTA injection, the roll ratio of the larvae was significantly reduced but larvae still did not reach the performance of the control fish. Tip links have been shown to regenerate within 8 to 24 hours^32^. Thus, the observed progressive recovery of postural control reflects the regeneration of the tip links, but may be slowed by residual BAPTA in the otic vesicle after the injection. We performed similar tests on buffer- and ferrofluid-injected fish, in order to disentangle the effect of the injection procedure from the ferrofluid itself on the vestibular system. Two hours after injection, we measured a roll ratio of 0.51 *±* 0.31 for the buffer injected fish and of 0.65 *±* 0.21 for the ferrofluid injected fish. Including the control group, we did not identify a significant difference between these three treatments (ctr, buffer, ferrofluid), neither at 2 h nor at 24 h or 48 h post injection (p = 0.456, 0.069 and 0.127, Kruska-Wallis test). Given the large inter-individual variability in postural control at this early developmental stage, we cannot exclude an effect on the function of the vestibular system from these data, but this effect should be mild and recoverable.

To test the immediate effect of the injection procedure on the mechanotransduction apparatus, we injected 4-Di-2-ASP, which fluorescently labels functional hair cells as it diffuses through the mechanotransduction channels when mechanotransduction is functional. Hair cells were clearly labelled after the injection (see inset Figure 2B). Thus, neither the injection procedure itself nor the ferrofluid appears to have a significant effect on the function of the vestibular organs.

Finally, we examined the kinematics of free swimming behavior in ferrofluid-injected fish compared to uninjected control larvae. We found that distributions of the inter-bout intervals and turning angles were not significantly different from control fish, while the average displacement per swim bout was only slightly but significantly increased (Figure S2 and Video S4). The various tests confirmed that the ferrofluid injection procedure had a very limited impact on the functionality of the vestibular system and on the locomotor behavior.

### Numerical simulations of the magnetic force exerted onto the magnetized otolith

To evaluate the impact of magnetic forces on the nanoparticle-covered otolith and its dependency on the magnet position relative to the larva, we resorted to numerical simulations (Figure 3). This approach revealed the existence of a range of magnet positions, within which the lateral force exerted onto the magnetized particle varied linearly with the lateral distance to the center of the magnet. The lateral force was maximal when the particle was located above the edge of the magnet, beyond which it decreased and eventually vanished. The maximal lateral force and the extent of the linear regime increased with magnet diameter, while the maximal lateral force decayed as z*^−^*^4^ as expected for a magnetic dipole, where z is the vertical distance to the magnet. These results suggest that the magnet should be placed as close as possible beneath the fish and that horizontal displacements should remain smaller than the radius of the magnet. Under these conditions, the force-displacement relationship is expected to be linear.

**Figure 3:**
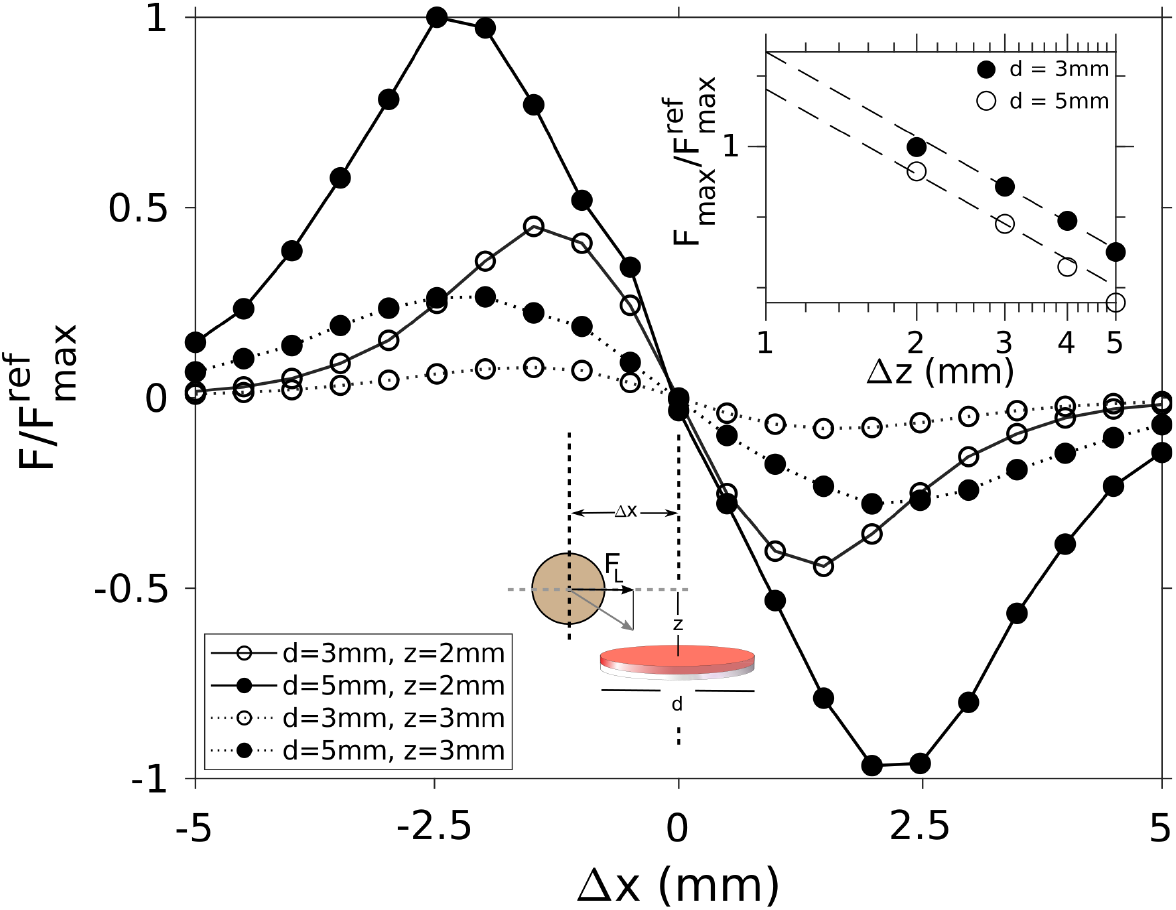
Finite element simulation of the lateral force, F*_L_*, exerted on a paramagnetic particle by a magnet as a function of the lateral distance (Δx) of the particle to the magnet’s center axis. Shown are force-displacement relationships for two magnet diameters, d, and two z-distances, z, between the magnet and the particle. The calculated forces were normalized by the maximum of the force-displacement relation, 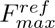, extracted from the configuration with the 5 mm in diameter magnet positioned at a z-distance of 2 mm. Inset: Dependency of the normalized maximal lateral magnetic force as function of the z-distance between the particle and the magnet, calculated for two magnet diameters (log-log scale plot). See also Figure S3.

To estimate the maximum force that can be imposed onto the otolith, we measured the velocity in water of an isolated utricular otolith (obtained after dissection of an injected larva) submitted to a comparable magnetic field as in the *in vivo* experiment. Taking into account the otolith diameter, that controls the drag force, we obtained an estimated force of *∼* 1 nN (see Star Methods). We can then compare this value to the gravitational forces exerted on the otolith *in vivo* when the head is pitch-tilted in space. The utricular otolith in fish is a calcium carbonate (aragonite) crystal with a density of 2.93 g.cm*^−^*^333^ and a diameter of *∼* 55 µm at the age of 6 dpf^10^. From these values, we estimated that under natural conditions, the maximal gravitational force experienced by the utricular otolith is *∼* 1.5 nN when the fish is rolled 90° (see Star Methods). The estimated magnetic and gravitational forces acting on the otolith are thus of the same order of magnitude, which *a posteriori* explains the capacity to drive large vestibular-like behavioral responses as described above. Our simulations can also be used to evaluate the number of nanoparticles attached to the otolith (for details, see Star Methods). A single particle experiences *∼* 0.007 fN of lateral force when placed at the edge of a 5 mm in diameter magnet positioned 2 mm beneath the particle. Therefore, approximately 0.6*·*10^8^ particles are required to produce a force of F*_g_*(15°) = 0.4 nN that would act on an otolith when the fish is rolled by 15°. This amount corresponds to approximately one monolayer of particles bound to the otolith (see Star Methods). This fine coating represents only 0.2 0 of the mass of the otolith and is thus unlikely to interfere with the vestibular function, in agreement with our observations.

### Brain-wide functional imaging during magnetic vestibular stimulation and behavioral monitoring

One of the assets of our stimulation technique is its low footprint, which facilitates a combination with any functional recording technique. Here, we used a setup that enables the application of controlled fictive vestibular stimuli in head-tethered larval zebrafish while recording the behavioral responses as well as the evoked brain-wide neuronal activity using light-sheet imaging (Figure 4A).

**Figure 4:**
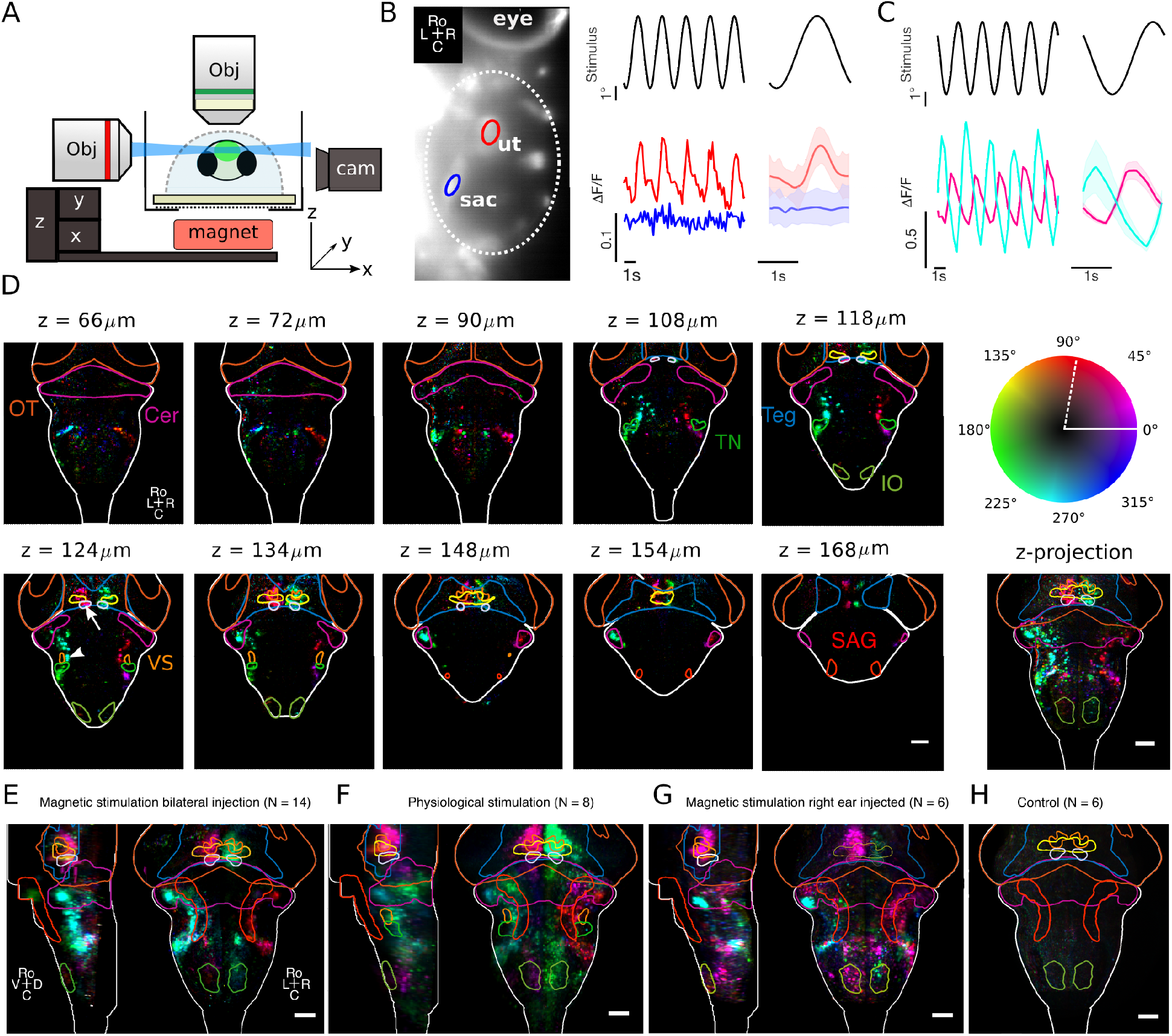
Brain-wide neuronal responses evoked by magnetic vestibular stimulation. Caption continues next page. **A** For functional imaging a light-sheet (blue) excites the fluorescence GCaMP6 sensor genetically encoded in the brain (green). The fluorescence is collected by an objective perpendicular to the light-sheet. **B** Calcium responses recorded in the otic vesicle from the utricle (red) and the saccule (blue) in response to a sinusoidal fictive roll stimulus (black). The trial-averaged response with std (N = 75 repetitions) is shown on the right. The ROIs from which the fluorescent signals were extracted are indicated with red and blue circles respectively. **C** Neuronal responses to the same stimulus as in **B** but measured in vestibulo-spinal neurons (white arrowhead in panel D) and extraocular motoneurons in the oculomotor nucleus (white arrow in panel **D** bottom left). **D** Selected layers of the phase map of the brain-wide response recorded in one fish. OT: optic tectum, Cer: cerebellum, Teg: tegmentum, nIII and nIV: oculomotor and trochlear nucleus, nMLF: nucleus of the medial longitudinal fascicle, SAG: statoacoustic ganglion, IO: inferior olive, TN: tangential nucleus, VS: vestibulo-spinal neurons. The color map indicates the phase of the neuronal response relative to the stimulus after correction for the phase delay introduced by the GCaMP6f calcium sensor Δϕ = arctan(*−*2πfτ_GCaMP6f_) with τ_GCaMP6f_ ≈ 1.6 s^37^. Without this correction the zero degree phase shift would be at the position indicated by the dashed white line. Bottom right: Maximum z-projection of the entire phase map shown of this fish. **E** Average phase map in response to bilateral fictive vestibular stimulation (N = 14 fish). **F** Average phase map recorded during natural vestibular stimulation with a rotating light-sheet microscope^6^. Note that the imaged volume largely overlaps with the volume imaged in E but was slightly shifted dorsal compared to the phase map shown in E. **G** Average phase map in response to unilateral (only right ear injected) fictive vestibular stimulation. **H** Control phase map recorded under the same conditions as E and H but without injected ferrofluid into the inner ear. Note that in D-H we do not show the rostral part of the phase maps where eye movements lead to brain deformations and thus to artifacts in the phase maps. Transgenic lines: *Tg(a-tubulin:Gal4-VP16; UAS:GCaMP7a)* panel B, *Tg(elavl3:H2B-GCaMP6f)* panel C-H. See also Video S5.

We first recorded the neuronal activity, evoked in mechanosensitive inner ear hair cells. In order to do so, we used the transgenic line α*-tubulin:Gal4-VP16;UAS:GCaMP7* ^34^, which we found to expresses the GCaMP7 calcium indicator in hair cells of the inner ear. Both inner ears of these fish were injected with ferrofluid, fish were embedded in agarose, and placed in the experimental setup one day after the injection. We generated a fictive vestibular roll-tilt stimulus by moving the magnet sinusoidally along the left-right body axis at 0.5 Hz and 2.5 mm amplitude. Hair cells in the anterior macula (AM, utricle) showed a modulation of the fluorescence signal, phase-coupled to the stimulus (Figure 4B). In contrast, hair cells in the posterior macula (PM), which are part of the saccule that senses vertical motion oscillations, showed no detectable response indicating that they were likely not stimulated. These observations are consistent with the anatomical orientation of the mechanosensitive axes of the two vestibular organs.

Next, we recorded the brain-wide neuronal dynamics elicited upon fictive vestibular stimulation using the pan-neuronal nuclear localized *Tg(elavl3:H2B-GCaMP6f)* transgenic line. This stimulus evoked neuronal activity throughout the brain (see Video S5). As an example, we show activity time traces recorded from vestibulo-spinal neurons and from the oculomotor nucleus nIII (Figure 4C). We quantified the brain-wide vestibular response pattern by computing a phase map as described in Migault *et al.*^6^. Briefly, we estimated, for each voxel (0.6×0.6×10µm), the amplitude and the phase relation of the evoked signal relative to the stimulus waveform. These two parameters were displayed in the form of a phase map, where color represents the relative phase of the neuronal response to the stimulus and intensity encodes the amplitude of the response. Hence, a phase shift of 0° applies to neurons whose activity is locked to the applied force, whereas a phase shift of 90° corresponds to neurons responding to the time-derivative of the force signal. Figure 4D shows the phase map for several selected layers and their z-projection, recorded in a single fish and registered on the Z-brain atlas^35^. The observed ocular motoneuron activity is consistent with the monitored compensatory eye movements. Active regions also include vestibulo-spinal neurons, the nucleus of the medial longitudinal fascicle, as well as hindbrain pre-motor neuronal populations involved in tail motion. The activity of cerebellar and inferior olivary neurons were also clearly modulated. The tangential nucleus appeared not in the phase maps because of a lack of labeling in the used transgenic line. The recorded response map was found to be stereotypic and reproducible for all injected fish, as shown by the sharpness of the average phase map, which combines the observations in 14 larvae (Figure 4E). This average phase map shows a close similarity with the brain-wide response recorded during natural vestibular stimulation using a rotating light-sheet microscope (Figure 4F and Figure 4 in Migault *et al.*^6^).

This new stimulation method offers, as optical tweezers, the opportunity to stimulate a single ear at a time, by injecting the ferrofluid only unilaterally. Figure 4G shows the average phase map for fish injected with ferrofluid into the right ear only. Interestingly, the response appears rather similar to that evoked by bilateral fictive vestibular stimulation, albeit with a relatively lower intensity. The antisymmetric activity in response to unilateral ear injection suggests an implication of commissural connections between both sides of the brain with a contralateral inhibitory connectivity in the hindbrain^36^.

We finally performed control experiments with non-injected fish. At the stimulation frequency, no signal was detectable in the average phase map (Figure 4H). This rules out the possibility that the recorded activity may in part reflect the visual stimulus caused by the moving magnet, which could have been possible as the blue (488 nm) laser forming the light sheet also illuminates the sample chamber.

## 3 DISCUSSION

Using iron particles to manipulate gravity sensation *in vivo* with magnetic fields to study balance and orientation has a long history. Already at the end of the 19th century, Kreid showed in crayfish (Decapoda, Crustacea) that the little stones that decapods collect from their environment to use as statoliths for gravity sensation could be replaced by ferrite grains. Placed in a magnetic field, animals then oriented their body relative to the position of the magnet. When the magnet was placed on top of the animal, they compensated by flipping on their backs. These experiments demonstrated that the small stones (statoliths) in the statocyst were used to detect gravity and accelerations by the decapods^38,39^. In 1978, Ozeki et al. carried out a similar study to characterize the response of the statocyst afferent neurons^40^. In contrast to decapods, vertebrates grow their statocyst in the inner ear, rendering experimental perturbations somewhat more difficult. Despite this challenge, we have developed a method that allows covering the otoliths of larval zebrafish with paramagnetic particles by injecting a ferrofluid into the developing inner ear. We demonstrated that subsequently, a magnet could apply forces to the utricular otolith *in vivo*, mimicking natural motion-like vestibular stimuli in immobilized animals. By using a small permanent magnet, we elicited robust motor responses that resembled those observed during natural vestibular stimulation, in both roll and pitch axis. While we used a custom-made ferrofluid for our experiments, similar ferrofluids, composed of negatively charged nanoparticles of similar diameter, are commercially available (see Star Methods section Ferrofluid), and can be used to evoke similar behavioral responses (see Figure S1 C).

Control experiments confirmed that the injection procedure did not damage the vestibular system and left the swimming behavior and postural control performance unaffected 24 hours after the injection. This robustness of the method reflects the minor interference of the injection procedure with the functionality of the system, but may in part be related to the capacity of non-mammalian hair cells for self-repair^32^. Hence, even if tip-links were damaged by the injection procedure, the vestibular apparatus is likely fully functional 24 - 48 h post injection. In addition to potential tip-link repair, non-mammalian ear hair cells can regenerate destroyed hair cell bundles^41^ and even entire hair cells with restored sensory function after cell death^42,43,44^. In the adult zebrafish utricle, the full regeneration of the utricular macula after induced damage takes about 13 days^45^. This is too slow to explain the observed high performance of the vestibular system after injections, leading to the conclusion that the injections did not cause substantial and functionally detrimental cell death.

The proposed mechanism underpinning the fictive stimulation is based on the irreversible binding of nanoparticles onto the surface of the otolith. This thin magnetized coating can then be acted upon using magnetic field gradients. We reported evidence of the effective magnetization of the otolith and that the corresponding magnetic force is in the same order of magnitude as the gravitational force imposed during macroscopic body rotation. In the present study, we only tested two different concentrations of injected ferrofluid, and the strongest response was obtained already at the lowest concentration. This observation suggests that a relatively small number of nanoparticles is sufficient to entirely cover the otolith with a compact monolayer. Any further particles are then likely to be repelled from the surface due to the electrostatic repulsion between the negatively charged nanoparticles. This coating was stable for at least four days after the injection, but likely even much longer.

A complementary mechanism may be at play that would rely on the magnet-induced motion of freely floating nanoparticles in the endolymphatic otic environment. The induced fluid motion would then impose a drag force onto the otolith. Given the nano-bead dimensions, the associated flow would be in the low Reynolds number regime and the particles are thus expected to reach their terminal velocity in less than a millisecond when placed in a field gradient. The resulting drag force, proportional to this particle velocity, would vary with the magnet position and could not be distinguished from the direct magnetic actuation on the otolith. However, an estimate of the particle terminal velocity results in a value of ≈ 0.1 µm/s (see Star Methods), which in turn yields a drag force orders of magnitude smaller than the force exerted by the particles attached to the otolith. This suggests that this second mechanism is probably negligible.

In zebrafish, the utricular otolith is spherical. For a spherical mass, the gravitational force grows with the radius cubed, while the magnetic force acting on a thin surface coating grows with the radius squared. One may thus anticipate that this magnetic actuation method should become relatively inefficient for larger animals (with larger otoliths). However, in most animals other than teleost fish, the otolithic membrane is covered with small carbon crystals called otoconia yielding a flat meshwork of extended mass. This leads to a much higher surface to volume ratio, which is more favorable for the actuation via surface-bound nanoparticles. Our method could thus work also in larger animals such as for instance *Xenopus* larvae, lampreys or even mammalian species provided that sufficiently strong magnetic fields and field gradients can be delivered. However, the interpretation of such experiments might be more difficult because different vestibular organs might be simultaneously acted upon. In this respect, the zebrafish larva appears to be a very convenient model system. In fact, a pilot study on an isolated *in vitro* preparation of *Xenopus* tadpoles at mid-larval stage, demonstrated that solutions of citrate-coated ferromagnetic nanoparticles can be reliably injected and distributed throughout the duct system of the inner ear. Repetitive displacement of a permanent point magnet above the transparent otic capsule in different directions, elicited faithful and robust eye movements also in these animals (see Video S6), which have a body size that is an order of magnitude larger than that of larval zebrafish. However, the fact that all the inner ear organs are functional at this developmental stage^46^, renders the accurate identification of the recruited vestibular organ(s) more difficult. In fact, the magnetic stimulation may drive multiple inner ear organs. Movement of the magnet approximately along the plane of the posterior semicircular canal produced movements of the ipsilateral eye in a corresponding oblique direction. Given these eye movements, we hypothesize that the posterior semicircular canal might in fact be actuated, since neuronal signals from this latter vestibular sensor represents the major activation pathway to superior oblique motoneurons to produce the observed eye movements^47^. This actuation is probably not mediated by flow induction, because the terminal drag velocity of the over-damped particles is very slow (see discussion above). However, we cannot exclude this possibility because the cupula, the mechanosensitive structure in the semicircular canals, is very sensitive. An alternative explanation is that the paramagnetic nanoparticles are bound directly to the cupula. Forces applied to the particles in a dynamic magnetic field gradient could then lead to a deflection of the cupula, mimicking the action of endolymph flow occurring under natural conditions, thus simulating vestibulo-ocular reflex activity. Furthermore, in our experiment, the point magnet probably does not activate the utricle, which is positioned more rostral, otherwise this would drive eye rotations in a different direction.

This magnetic actuation method was implemented to fictively stimulate slow rollor pitch tilts of the fish body. However, the same approach could be used to mimic translational accelerations experienced by the larvae during free swimming. Larval zebrafish swimming patterns consist of discrete swim bouts that last for 100 *−* 200 ms interspersed by *∼* 1 s-long resting periods. Owing to the Einstein principle, otoliths are actuated during these transient linear accelerations: forward acceleration of the animal produces a backwards pointing force on the otolith while a deceleration corresponds to a forward pointing force. The reported peak acceleration during a bout is in the range of 0.3 *−* 2 m/s^2^ = 0.03 *−* 0.2 g^48,49^, which corresponds to a force on the otolith in the range of 50 *−* 300 pN, i.e., within the accessible range of our instrument. To mimic acceleration forces encountered during a swim bout, the magnet has to be moved by 0.3 *−* 2 mm in about 50 ms, which is also compatible with the performance of our mechanical stages. Our system can thus be used to emulate vestibular signals associated with fictive self motion in head-fixed animals. It could therefore be included into closed-loop virtual reality assays, mitigating sensory mismatch and enhancing the quality of virtual environments. This opens new possibilities to study sensorimotor processing.

Unlike other approaches, such as optical tweezers, the reported method could potentially be implemented in freely swimming configurations as well. A large scale magnetic field gradient could be used to create a sufficiently large force onto the magnetized otolith coating to counteract the gravitational force acting on it. One could thus create a zero gravity condition or mimic inverted gravity and study how fish adapt to and learn to cope with this change of physical parameters. As only injected fish will be sensitive to the applied magnetic field gradients, social behavior experiments can be envisioned to study how conspecific fish react to behavioral changes of a single fish when the latter experiences a perturbation of the vestibular sensation. But one may even go beyond and investigate how animals can learn to use this novel sensation of magnetic field gradients, e.g., for navigational strategies.

We have demonstrated that our vestibular stimulation method is compatible with simultaneous whole-brain functional imaging using light-sheet microscopy. In response to the fictive vestibular stimulus, we observed consistent neuronal activity in the vestibular nucleus and in downstream nuclei throughout the brain. The evoked neuronal response map in bilaterally injected fish was comparable to the one that was obtained during actual vestibular motion stimulation with a rotating light-sheet microscope. The recorded average phase maps during unilateral stimulation suggest the presence of a pronounced commissural inhibition between the two vestibular nuclei, typically conserved in all vertebrates^36^. This is consistent with the results obtained by unilateral utricular stimulation using optical tweezers [7]. Sinusoidal magnetic stimulation of the right ear shows that pulling the right otolith laterally activates neurons located in the right vestibular nucleus and downstream regions such as the ipsilateral vestibular cerebellum, and on the contralateral side oculomotor motoneurons, neurons in the nMLF as well as hindbrain neuronal populations probably projecting to the spinal cord. This activity pattern and profile is consistent with the highly conserved axonal projections from the vestibular nucleus to these brain regions^50,51^. This activation pattern has a mirror-symmetric counterpart with a mean activity that is 180 degrees phase-shifted, and thus exhibits a mean activity that is minimal when the mirror-symmetric neuronal correlate is maximally active. This suggests that the vestibular nucleus inhibits the contralateral vestibular nucleus, which leads to a reduced activity in downstream nuclei. The latter result is consistent with the description of inhibitory commissural projections in cats between vestibular neurons of the utricular pathway^52^, which are thought to contribute to the sensitivity of vestibular neurons through a disinhibition^52^. In larval zebrafish, commissural projections have been described as originating from the tangential vestibular nucleus^9^, with a likely inhibitory function as evidenced by our results.

In summary, our magnet-based vestibular stimulation method is inexpensive, easy to implement, and can be developed as an add-on device for existing microscopes and visual virtual reality setups. Since the magnet is small and operates beneath the fish, the whole experimental chamber is accessible for all types of microscopes, optogenetic tools, electrophysiological setups, other sensory stimulation methods or behavioral monitoring. Accordingly, our method uniquely expands the toolbox of widely accessible sensory stimulation methods for zebrafish systems neuroscience but also for neuroscientific studies in other species.

## ACKNOWLEDGEMENTS

We thank Christine Ménager and Aude Michel Tourgis (Sorbonne Université, Laboratoire PHENIX, CNRS UMR 8234) who kindly provided the ferrofluid. We thank the IBPS fish facility staff for the fish maintenance, in particular Stéphane Tronche and Alex Bois. We thank Misha Ahrens and Teresa Nicolson for providing transgenic fish lines. We are grateful to Carounagarane Dore for his contribution to the design of the experimental setup. We thank Claire Wyart and Marcus Ghosh for their comments on the manuscript. This project has received funding from the European Research Council (ERC) under the European Union’s Horizon 2020 research innovation program grant agreement number 715980, and was partially funded by the CNRS, Sorbonne Université, and by the German Science Foundation through the collaborative research center 870 (CRC 870). H.M., G.M., and P.T. had a PhD fellowship from the Doctoral School in Physics, Ile de France (EDPIF). G.L.G. had a PhD fellowship from the Systems Biology Network of Sorbonne Université.

## AUTHORS CONTRIBUTION

N.B., G.D. and V.B designed the project. N.B., H.M., G.M. and P.T. performed the zebrafish experiments. N.B. T. Panier, T. Pujol, G.M, and V.B. built the experimental setup. T.Pujol performed the finite element simulations. N.B., H.M., G.M., G.L.G., P.T., G.D., V.B. analyzed the data, N.D. and H.S. performed the *Xenopus* experiment. N.B., G.D., V.B wrote the manuscript with input from all the authors.

## STAR*METHODS

## RESOURCE AVAILABILITY

### Lead contact

Further information and requests for resources and reagents should be directed to and will be fulfilled by the lead contact, Volker Bormuth.

### Materials Availability

All the fish lines used in this study are listed on the key resource table. This study did not produce new transgenic lines. All the lines used are already published, and available upon request. The custom-made ferrofluid used in this manuscript can be replaced by a commercially available ferrofluid. We tested this ferrofluid and it works equally well. This ferrofluid is available directly from the website: https://ferrofluid.ferrotec.com/products/ferrofluid-emg/water/emg-304/. This study did not generate further new unique reagents.

### Data and code availability

Data and code are available via this link: https://psilo.sorbonne-universite.fr/index.php/s/Beiza2023. Any additional information and large raw data videos will be shared by the lead contact upon request.

## EXPERIMENTAL MODEL AND SUBJECT DETAILS

### Animal husbandry

All experiments were performed on 5-9 dpf larvae. The sex of the animals was not yet determined at this age, and was therefore not reported. Adult fish were maintained at 28°C in system water (pH 7-7.5 and conductivity between 300 and 350 µS) in the fish facility of the Institut de Biologie Paris-Seine. Eggs were collected in the morning and then kept in a Petri dish with E3 at 28°C under a 14h/10h light/dark cycle. Larvae were fed with rotifers from 5 dpf on. Calcium imaging experiments were carried out in two different transgenic lines: *elavl3:H2B-GCaMP6f* ^53^ (kindly provided by Misha Arhens) and α*-tubulin:Gal4-VP16; UAS:GCaMP7* ^34^ (kindly provided by Teresa Nicolson), both in nacre background. Homozygous nacre mutants lack melanophores, which makes them more suitable for imaging. Experiments were approved by Le Comité d’Éthique pour l’Expérimentation Animale Charles Darwin C2EA-05 (02601.01 and #32423-202107121527185 v3).

## METHOD DETAILS

### Ferrofluid solution

The ferrofluid, a suspension of γ-Fe_2_O_3_ iron oxide nanoparticles, was produced by Christine Ménager and Aude Michel Tourgis (Sorbonne Université, Laboratoire PHENIX, CNRS UMR 8234) following the protocol described by Massart et al.^25^ and kindly provided to us for our experiments. The hydrodynamic diameter measured by dynamic light scattering (DLS) was 22 nm with a polydispersity index of 0.15. This corresponds to a physical diameter of 11 nm, usually measured by TEM after drying the sample. The particles were dispersed in water and stabilized with citrate molecules at pH 7 to prevent agglomeration. Similar water-soluble particles are commercially available from ferrotec.com, such as the EMG 304 product, with the same diameter. These molecules are also negatively charged but not via covalently bound citric-acids as in our study but via absorption of a negatively charged polymer. We tested also this ferrofluid solution and it worked equaly well.

### Ear injections

Either ferrofluid, BAPTA or 4-Di-2-ASP were injected into the inner ear with a glass micropipette held by a micromanipulator (Narishige MN-153) using a pneumatic Pico-pump (World Precision Instruments PV830). Capillaries (1 mm outer diameter, Warner Instruments GC100F-10) were pulled to obtain fine tip micropipettes (tip diameter = 1 to 2 µm) using a Narishige PC-100 puller with the following parameters: 2 steps, Heater N°1 = 52.4, Heater N°2 = 55.7, position 2 mm, 2 heavy and 1 light weights. Micropipettes were loaded with 2 µL of ferrofluid diluted in buffer (NaCl 0.178 M, sodium citrate 0.023 M, HEPES 0.01 M). Injections were performed at 5 dpf. On top of a microscope glass slide, larvae were mounted dorsal side up in 2 % low melting point agarose. Using a small piece of metal as support, the slide was tilted by 45 degrees to access the fish’s left ear. For the ferrofluid, 3 pulses (10 psi for 500 ms) were injected into the otic vesicle, corresponding to a total volume of 1,2 nL. After the left ear was injected, the glass slide was tilted onto the other side to inject the right ear. For injection of BAPTA and 4-Di-2-ASP the protocol was the same. We used 50 mM BAPTA dissolved in extracellular solution containing (in mM) 134 NaCl, 2.9 KCl, 1.2 MgCl_2_, 2.1 CaCl_2_, 10 HEPES, and 10 glucose, at 290 mOsm, adjusted to a pH of 7.8 with NaOH. For 4-Di-2-ASP, a 50 mM solution of diluted E3 medium containing 1% ethanol was injected. After the injections, the larvae were freed from the agarose with fine-tipped forceps (Dumont n°5) and maintained in E3 medium until the start of the experiments.

### Roll ratio essay

The larvae were placed in a 5 cm Petri dish positioned under a high magnification objective and recorded at 300 fps. Approaching the larvae with a fin glass tip or by inducing a vibration evoked a startle response. Each larva was subjected to five trials. The roll behavior was assessed for each trial. The roll ratio was calculated as the number of trials the animal rolled during an escape divided by the number of trials the animal attempted an escape^30^.

### Sample preparation

24 hours after ferrofluid injection, larvae were mounted in 2 % low melting point agarose dorsal side-up on top of a small acrylic holder (1mm thick). Then, the holder was placed inside an acrylic chamber filled with E3. For behavioral experiments, the agarose was removed from the eyes and tail using a micro knife (FST Micro Knife – Plastic Handle/22.5° Cutting Angle).

For the neuronal recordings under physiological vestibular stimulation, the fish were paralyzed before being mounted in agarose by bathing them for 2-5 min in a solution of 1 mg/mL α-bungarotoxin (Thermofisher Scientific) in E3 medium. We then transferred them into pure E3 medium and waited ≈30 min to check for absence of motor activity and normal heart beating.

### The setup

We built a platform for the magnetic sitmulation with two motorized stages (Physik Instrumente, V-408 PIMag Linear Stage) to precisely control the magnet position in X and Y and hence the fictive vestibular stimulation. A third manual stage allowed to position the magnet beneath the fish as close as possible in the vertical plane in order to maximize the accessible range of force. For the experiments shown in Figure 1 and 4 we used a magnet 5 mm in diameter and 3 mm in height.

Injected fish were mounted on a 1 mm thick transparent acrylic holder. The fish were held in a drop of low melting point agarose at approximately 1 mm above the surface. The holder was then placed in the sample chamber filled with embryonic medium E3. The bottom of the sample chamber was formed by a 220µm thick glass coverslip.

The magnetic stimulation unit was mounted on an imaging setup built around a microscope frame from Scientifica (Scientifica Slicescope Pro) fitted with an Olympus BX-URA fluorescence illuminator and a custom light-sheet forming unit adapted from Migault et al.^6^. Functional imaging was performed with a Leica HC FLUOTAR L 25x/0,95 W VISIR objective and a Hamamatsu Orca-Flash4.0 V3. Images were recorded with HCImage software (Hamamatsu) and the light-sheet was controlled with a custom application written in Matlab (MathWorks). Pixel size in the images were 0.58µm.

Top view, behavioral tail recordings used the microscope’s light path and camera with a Nikon CFI Achro 4x objective. Side and front view behavioral recordings used separate systems of Point Grey cameras (BFLY-U3-05S2M-CS) with Navitar Precise Eye objectives (1-61450 and 1-61449).

Behavioral and neuronal responses to physiological vestibular stimulation were recorded on our rotating light-sheet microscope [6]. Pixel size corresponded to 1.2µm for in the neural recordings.

All neuronal data were recorded at a volumetric acquisition rate of 4Hz. For each stack, 25 layers separated by 10µm were recorded. The total length of each recording was 5min.

### Behavioral protocol

To simulate a roll-like motion, we moved the magnet along the transverse axis, starting from the center and extending 2.5 mm towards each side of the fish. To simulate a pitch-like motion, we moved the magnet along the longitudinal axis, using the same amplitude. The stimulation frequency in both cases was 0.5 Hz. The step stimulus was 2.5s to each side with a 2.5s dwell time at zero before each step. Eye movement responses were recorded at a rate between 15 and 30 frames per second.

### Force generation mechanism

#### Force of a ferrofluid particle in a magnetic field gradient

The ferrofluid particles are so small that nanoparticles consist only of a single magnetic domain, giving the particle a giant magnetic moment. In the absence of an external magnetic field, the direction of this moment changes randomly depending on the temperature. The average magnetisation is zero and the particle is in a superparamagnetic state. In an external magnetic field, the giant magnetic moment becomes progressively aligned against the thermal agitation, and the average net magnetization increases. The macroscopic magnetization of a ferrofluid particle or of a ferrofluid droplet is characterized by the macroscopic magnetic moment, 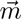, which depends on the volume V of the particle or of a ferrofluid droplet, and the external field B. In a weak magnetic field the macroscopic magnetization is given by

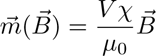

with χ the magnetic susceptibility and µ_0_ the vacuum permeability. And the force exerted on the droplet reads

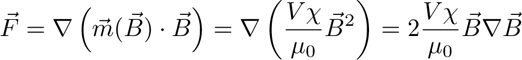

In a strong field that saturates the magnetization, the force exerted on the droplet is

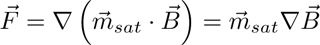

For details see e.g. Baster 1987^54^.

#### Finite element simulations

We used Comsol Multiphysics to calculate the magnetic force applied by the magnet to the ferrofluid. Lateral force-displacement curves were calculated for cylindrical magnets of different diameters and z-distance to a spherical droplet of the ferrofluid with a diameter of 200 µm. The spherical droplet was considered perfectly rigid. The droplet volume was chosen arbitrarily. The force, acting on the droplet, depends linearly on the volume of the droplet. Therefore, uncertainty with respect to the droplet volume will change the maximum force reached, but not the linear dependence of force on the magnet position. The relationship between magnetic flux density and magnetic field strength (B-H curve) is defined for the ferrofluid by a magnetization curve (Figure S3). The magnetic flux density is fixed for the magnets. For the simulations, we started with a mesh size of 500 µm and then iteratively reduced the mesh size until the results converged. Parametric sweeps were realized for different distances and diameters.

The simulation gives a maximal lateral force of F*_D_*_=200_ *_µm_* = 4 *·* 10*^−^*^4^N exerted on the ferrofluid droplet with a diameter of D = 200 µm placed 2 mm above a 5 mm in diameter magnet. As the force depends linearly on the volume, we estimated that the force exerted on a single nanoparticle with a diameter of D*_p_* = 11 nm was F*_p_* = 0.007 fN. Analytical equations to calculate the force extension curve as well as how the forces scale with magnet size have also been developed and can be found here^55^.

#### Gravitational force F*_g_*exerted onto the otolith during roll motion

Under natural conditions, when the fish is rolled along the rostro-caudal body axis, gravity acts on the otoliths pulling them along the left-right body axis. The magnitude of this lateral component of the gravitational force F*_g_* depends on the roll angle

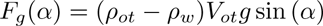

with the density of the otolith ρ*_ot_* = 2.93 g cm*^−^*^333^, the density of water ρ*_w_* = 1 g cm*^−^*^3^, the otolith volume 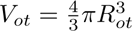, the otolith radius R*_ot_* = 27 nm, the gravitational acceleration g = 9.81 m s*^−^*^2^ and the angle α by which the animal is rolled relative to its dorsal side-up position.

At α = 90° the lateral force on the otolith is maximal with

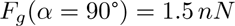

#### Estimation of the number of nanoparticles bound to the otolith

Because the mean behavioral response in the fictive roll motion experiments compares to the mean evoked response when rolling fish with a sinusoidally modulated excursion of *±*15°, we can estimate that we exerted *in vivo* with our experimental parameters in average a maximum force of < F*_max_* >= 1.5 nN *·* sin(15°) = 0.4 nN on the otolith when displacing the magnet 2.5 mm.

Given the force that the magnet exerts on a single particle of the ferrofluid suspension F*_p_* = 0.007 fN, we can estimate the number of particle bound to the otolith

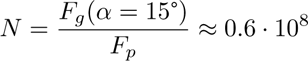

A monolayer of particles on the otolith surface corresponds to approximately

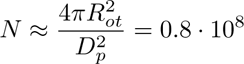

particles with D*_p_* = 11nm the diameter of the nano particles. Thus, we estimate that *∼* 1 monolayer of particles has bound to the otolith.

However, due to the small diameter of the particles, the mass change of the otolith is negligible with

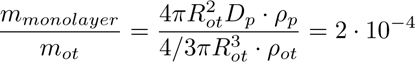

with ρ*_p_* = 4.8 g cm*^−^*^3^ the density of the ferrofuid particles (maghemite, γ *−* Fe_2_O_3_) (www.matweb.com), and ρ*_ot_* = 2.93 g cm*^−^*^3^ the density of the otolith^33^.

#### Drag force on an otolith pulled through water

To estimate the maximum force that can be delivered to the otolith, we measured the velocity in water of an isolated utricular otolith (obtained after dissection of an injected larva) submitted to a comparable magnetic field as in the *in vivo* experiment (see Figure S1A). Taking into account the otolith diameter that controls the drag force, we obtained, using Stokes’ law, an estimated force of

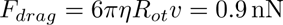

with η the viscosity of water R*_ot_* = 27.5 µm, the radius of the otolith and v = 1.75 mm/s the measured speed at which the otolith was dragged by the magnet through the aqueous solution. This estimated force is of the same order of magnitude as the estimated force of gravity acting on the otolith during a roll stimulus.

#### Time constant at which a particle reaches its terminal velocity when accelerated by a constant force in a viscous solution

Freely floating particles in the inner ear will be accelerated by the magnet. However, due to the interaction with the surrounding water molecules they will reach a terminal velocity after a characteristic time^56^

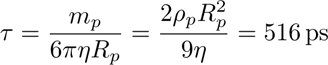

with the particle mass m*_p_*, the hydrodynamic particle radius R*_p_*= 22 nm, and the viscosity of water η. We calculated a terminal velocity of

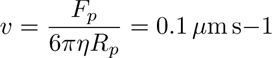

with the estimated maximal force F*_p_* = 0.007 fN exerted on a ferrofluid nanoparticle placed 2 mm over the edge of our 5 mm in diameter magnet.

#### Applicability to other animal models

Although it is difficult to precisely predict the intensity of the magnetic force one could deliver, e.g., in rodents with this technique, we can offer some basic scaling evaluation.

What is relevant in our experiment is the ratio of the magnetic force, F*_m_*, applied to the ear stone, and the gravitational force acting on it, F*_g_*. This indeed sets the extent to which we are able to mimic the effect of gravity. If we consider magnets of similar aspect ratio, but with different dimensions (defined by a scaling length k), then this ratio scales as:

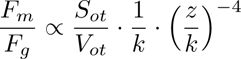

where z is the distance between the otolith and the surface of the magnet, S*_ot_* and V*_ot_* are the surface area and the volume of the otolith, respectively. The term S*_ot_*/V*_ot_* reflects the fact that the magnetic force varies linearly with the number of bound nanoparticles, which we assume to be proportional to the surface of the ear stone, whereas the gravitational force is proportional to the otolith volume. The second term of the expression derives from the dependence of the magnetic field produced by a cylindrical magnet with its dimension and the distance z to its surface (see Agas et al. 2008^55^).

In teleost fish that grow a spherical ear stones the ratio S*_ot_*/V*_ot_* is of the order of 1/D*_ot_* making our method less favorable for older fish that have otoliths larger in diameter.

In contrast, in mice the otolithic membrane is covered with small carbon crystals called otoconia yielding a flat meshwork of extended mass. The height, h, of this meshwork is several tens of micrometers^57^ so of the order of the larval zebrafish otolith diameter, D*_ot_*, and it extends over a disk of diameter 500 µm. As a consequence, one expects the ratio S*_ot_*/V*_ot_* = S*_ot_*/(S*_ot_ ·* h) = 1/h ≈ 1/D*_ot_*. So the ratio of S*_ot_*/V*_ot_* in mice is thus relatively similar to the one in larval zebrafish, for which we have demonstrated our method.

With larval zebrafish, we succeeded in obtaining a strong vestibular response by placing a cylindrical magnet (5 mm x 3 mm) at a distance of 2 mm from the utricle. The head of a standard laboratory mouse (houses mouse) has a skull height of approximately 10 mm. Scaling up the magnet by a factor k = 10 (results in a magnet dimension of 5 cm x 3 cm) and placing it at 10 mm distance from the otolith (i.e., in proximity to the head) would in principle yield a similar magnetic to gravitational force ratio as in our larval zebrafish experiment.

This simple scaling argument thus suggests that the same method should be applicable in rodents.

## QUANTIFICATION AND STATISTICAL ANALYSIS

### Free swimming control

We analyzed the free swimming behavior of ferrofluid-injected (N = 7) and control (N = 11) fish. Larvae were injected at 5 dpf. 24 hours later, they were placed in a Petri dish to record the swimming behavior for 1 hour at 30 fps. Videos were tracked using the FastTrack program^58^. Individual fish were not tracked through the entire video, but were split into wall-to-wall trajectories. For each trajectory, discrete swim bouts were detected when the instantaneous swim speed exceeded two times the total variance of the speed. Putative bouts were then filtered using a distance criterion (bouts with a linear displacement – measured in a time window of *±*0.5 s centered on the bout velocity peak – less than 0.3 mm or greater than 18 mm were rejected) and on a temporal criterion (bouts occurring within 0.4 s after a bout were rejected). Bout onset was defined at 80 ms before the velocity peak. From the positions, time and body angles before and after an event, the inter-bout interval, displacement, and turn angles associated with each bout (N*_control_* = 4184, N*_injected_* = 1039) were computed. These values were then averaged over the trajectories (N*_control_* = 257, N*_injected_* = 64) and the trajectory means were plotted as boxplots. Outliers are shown as blue circles. The mean square displacement (MSD) was calculated using the MATLAB package msdanalyzer^59^. The x,y-sequences were pooled by condition (control and injected), an MSD was calculated for each sequence. In Figure S2D the the ensemble mean is presented along with the standard error of the mean.

### Statistical Analysis

Sample size was different for each test and indicated in every figure. Data analysis and generation of plots was performed using the MATLAB functions *boxplot, adtest, kruskalwallis, signrank, anova1* and *multicompare*. All the boxplots show as center and dispersion measures the median and the IQR. Significance was defined as p-values smaller than 0.05.

### Behavioural analysis

Eye movements in response to magnetic vestibular stimulation at 0.5Hz were extracted online over time using a template matching algorithm implemented in LABVIEW (National Instruments). The peak-to-peak angular response amplitude was manually extracted from the first few cycles of the recorded response. When the template matching algorithm was unable to accurately track eye movement and introduced a lot of noise, the eye angle was manually extracted from the recorded video using Fiji. Eye movements in response to the physiological sinusoidal rolling stimulus at 0.2 Hz were recorded for 2 min. The recorded images were averaged offline by calculating the median over the stimulation cycles. To determine the eye angle in the averaged images at different times during the stimulation cycle, 4 landmarks were placed on the eye and two intersecting segments were formed. We followed the orientation of the segments over time during the stimulation cycle, which allowed us to determine the angular change in eye rotation between the maximum and minimum of the stimulation cycle. The peak-to-peak angular response was then defined as the average of the measurements obtained with the two segments. The tail movement in response to the magnetic vestibular stimulation was recorded at 50 frames per second. Using Matlab, we segmented the tail in ten segments and determined the tail angle as the difference between the angle of the first (most rostral) segment and the fourth segment. This reduced the contribution of transiently evoked swims to the measured tail angle and allowed us to extract the slow response locked to the stimulus frequency.

### Calcium imaging analysis

Drift correction, calcium transient (ΔF/F) extraction, and calculation of the phase maps were performed offline using MATLAB, according to the workflow previously reported^6^. XY drifts were corrected by registering a user defined subpart of each image of the stack with the corresponding subpart in the first image. Phase maps were established by calculating a Fourier transformation of the fluorescence time trace of each pixel and estimating the amplitude A and phase Φ at the stimulation frequency. A baseline A*_b_*was estimated as the mean amplitude over a frequency window encompassing 20 points to the left and right of the peak, with windows starting three points from the peak. We then computed the normalized response, equivalent to a signal-to-noise ratio, (A*−*A*_b_*)/A*_b_*. To compute the actual phase shift of the fluorescent response to the stimulus, we took into account that the control signal of the vestibular stimulus was a cosine phase-shifted by ϕ*_stim_* = *−*π/2 for the magnetic setup and ϕ*_stim_* = π for the rotating light-sheet microscope. Furthermore, we accounted for a synchronization delay between our stimulus control signal and the onset of the image acquisition of τ = 226 ms for the magnetic system and τ =302.2 ms for the rotating light-sheet microscope. This delay corresponded to an additional phase offset of ϕ*_offset_* = 0.6174 for the magnetic system and ϕ*_offset_* = −0.9494 for the rotating microscope at the stimulation frequency of 0.5 Hz. Accordingly, we computed the phase of the fluorescent response relative to the stimulus as ϕ_fluo_ = Φ + ϕ*_stim_* + Φ_offset_. To estimate the phase of the neuronal response relative to the stimulus, we subtracted the phase shift introduced by the calcium sensor (see [6]) ϕ_neuro_ = ϕ*_fluo_ −* Φ_GCaMP6f_, which we estimated to be Φ_GCaMP6f_ ≈ arctan(*−*2πfτ*_GCaMP_* _6_*_f_*) = *−*1.374rad = *−*78.74°. We estimate the nuclear GCaMP6f decay time constant τ = 1.6s^37^ using blind deconvolution for spike inference from fluorescence recordings^60^ applied to brain-wide recorded spontaneous activity.

We estimated the relative variations of the fluorescence intensity, ΔF/F, shown in Figures 4B-C, with respect to the baseline signal as ΔF/F = (F(t) - baseline)/(baseline - background). The background was neglected in B and estimated in C from the average intensity of pixels outside the brain. The baseline fluorescence signal was estimated for each ROI by a running 10th percentile estimation of the fluorescence time signal in a sliding window of 50 s.

### Registration onto the Z-Brain atlas

We used the Computational Morphometry ToolKit CMTK (http://www.nitrc.org/projects/cmtk/) to compute for every fish the morphing transformation from the average brain stack (anatomical stack) to the Elavl3:H2B-RFP stack of the Z-Brain atlas^35^. This allowed mapping the functional data onto the Z-Brain Viewer, to overlay the region outlines, and to calculate averages across animals. Automatic registration was not always satisfactory. To improve the mapping, we manually rotated along the pitch axis mapped stacks by 3 degrees. We computed first the affine transformation, which we used then as initialization to compute the warp transformation between the two stacks. The used commands and options are listed in table 1.

## KEY RESOURCES TABLE

**Table 1:** CMTK commands and options.

| Tool | Options | Description |
| --- | --- | --- |
| cmtk registration | -Initxlate<br>-dofs 6,9,12<br>-sampling 3<br>-coarsest 25<br>-omit-original-data<br>-accuracy 3<br>-exploration 25.6 | Calculate affine transformation |
| cmtk warp | -v<br>--fast<br>-grid-spacing 40<br>-refine 2<br>-jacobian-weight 0.001<br>-coarsest 6.4<br>-sampling 3.2<br>-accuracy 3.2<br>-omit-original-data | Use affine transformation as initialization |
| reformatx |  | Apply transformation to other stacks |

**Table 2:** Key resource table.

| REAGENT OR RESOURCES | SOURCE | IDENTIFIER |
| --- | --- | --- |
| **Chemicals** |  |  |
| Low melting point agarose | Sigma-Aldrich | A9414-50G |
| Tricaine | Sigma-Aldrich | E10521-10G |
| α-bungarotoxin | ThermoFisher Scientific | B-1601 |
| Ferrofluid | custom-made | Massart et al., 1995<sup>25</sup> |
| Ferrofluid | ferrotec | https://ferrofluid.ferrotec.com/products/ferrofluid-emg/water/emg-304/ |
| BAPTA | Sigma-Aldrich | 14510-100MG-F |
| 4-Di-2-Asp | Sigma-Aldrich | D3418-500MG |
| Ultrapure Low Melting Point Agarose | Invitrogen | 16520050 |
| **Experimental Models: Organisms/Strains** |  |  |
| Tg(a-tubulin:Gal4-VP16 ;UAS:GCaMP7a) |  | Koster et al., 2001<sup>34</sup> |
| Tg(elav3-H2B:GCaMP6f) |  | Vladimirov et al., 2014<sup>53</sup> |
| **Software and Algorithms** |  |  |
| Matlab | The MathWorks | https://www.mathworks.com/products.html |
| Labview | National Instruments | https://www.ni.com |
| CMTK | Rohlfing and Maurer, 2003<sup>61</sup> | https://www.nitrc.org/projects/cmtk/ |
| FIJI | (ImageJ) NIH | https://fiji.sc/ |
| Z-Brain atlas | Randlett et al., 2015<sup>35</sup> | https://zebrafishexplorer.zib.de/home/ |
| Comsol Multiphysics | Comsol | https://www.comsol.com/ |
| **Other** |  |  |
| Pneumatic PicoPump | World Precision Instruments | SYS-PV830 |
| Glass capillaries to pull micropipettes | Warner Instruments | GC100F-10 |
| Micropipette puller | Narishige | PC-100 |
| Motorized stages | Physik Instrumente PI | PIMag® Linear Stage: V-408.132020, V-408.232020 |
| Behavior tracking: Camera | Point Grey | BFLY-U3-05S2M-CS |
| Behavior tracking: Objective | Navitar | 1-61449 |
| Behavior tracking: 2x Adaptor | Navitar | 1-61450 |
| Magnet (D=5 mm, thickness 1 mm, 3 magnets stacked) | RS Components | N837RS |
| Micro knife 22,5° cutting angle | Fine Science Tools | 10316-14 |

## 4 VIDEO LEGENDS

### Video S1

#### Behavioral responses to fictive magnetic vestibular stimulation with a hand-held magnet

**Part I:** Behavioral response to fictive vestibular roll stimulation recorded by top view monitoring. The fish is tethered with agarose that was removed around the tail and eyes to allow free movements. The video was obtained with a stereo microscope immediately after ferrofluid injection. The magnet was hand-held and moved beneath the fish along the left-right body axis to mimic a vestibular roll stimulus.

**Part II:** Same fish as in Part I, with the magnet moved along the rostro-caudal body axis to mimic a pitch-tilt stimulus (nose-up, nose-down). Eyes and tail perform marked compensatory movements in response to these fictive vestibular stimuli. Related to Figure 1.

### Video S2

#### Behavioral responses to fictive vestibular roll- and pitch-tilt stimuli generated and recorded in an automatized setup

Ferrofluid was bilaterally injected at 5 dpf and videos were recorded the next day at 6 dpf. The magnet was moved sinusoidally at 0.5 Hz and with 2.5 mm amplitude.

**Part I:** Front view of evoked eye movements in response to a fictive roll stimulus. The eye rotates only along the roll axis and not along the pitch-tilt axis, demonstrating the absence of cross-talk between the two stimulation axes.

**Part II:** Side view of evoked eye movements in response to a fictive pitch-tilt stimulus.

**Part III:** Top view of evoked tail movements in response to a fictive roll stimulus. Note that the magnet created a shadow under the fish, which allows us to see the correlation between magnet displacement and tail movement. The tail performs a tonic tail pivoting movement to the contralateral side of the magnet movement, as well as discrete swim bouts. This is consistent with the observation by Favre-Bulle et al. [10] using optical tweezers to stimulate the vestibular system. When averaging over many stimulation cycles, the discrete responses average out.

**Part IV:** A bilaterally injected fish is shown in front view in two conditions. The left recording shows the fish in the absence of a vestibular stimulus. The eye orientation does not show a static bias, nor did we observe a smooth drift of the eyes that could potentially indicate a malfunction of the vestibular system. The right recording shows the fish responding to a physiological vestibular roll stimulus with a sinusoidal waveform of 0.5 Hz and an amplitude of *±*15°. Related to Figure 1.

### Video S3

#### Impaired postural control along the roll axis after BAPTA injection compared to wild-type control fish with an intact vestibular system

**Part I:** The video shows a fish two hours after bilateral injection of the calcium chelator BAPTA. The fish was placed freely in a Petri dish filled with embryonic medium E3 and was monitored from the top. A startle response was elicited by touching the fish with a fine glass tip. During the startle response and also during successive swimming bouts, the fish lost its dorsal-up posture and rolled around its longitudinal body axis. **Part II:** Recording of a wild-type control fish under the same conditions as in Part I. Throughout the sequence and including the startle response, the fish maintained its dorsal side-up posture. Related to Figure 2.

### Video S4

#### Free swimming behavior of fish, injected bilaterally with ferrofluid, compared to wild-type control fish

**Part I:** Wild-type fish (N = 10) swimming freely in a Petri dish (Recording frame rate = 70 fps)..

**Part II:** Fish after bilateral injection of ferrofluid into the inner ears (N = 7), swimming freely in a Petri dish (Recording frame rate = 70 fps). Related to Figure 2.

### Video S5

#### Brain-wide neuronal responses evoked by a fictive sinusoidal roll stimulus

The video shows in a loop the neuronal response averaged over 40 stimulus cycles. Six sections of the brain are shown.

*Experimental parameters*: Ferrofluid was injected bilaterally at 5 dpf into a fish of the Tg(elav3-H2B:GCaMP6f) transgenic line. The video was recorded at 6 dpf. The magnet was moved sinusoidally at a frequency of 0.5 Hz and with an amplitude of 2.5 mm. Indicated z positions are measured relative to the first imaged layer. Related to Figure 4.

### Video S6

#### Pilot study in a *Xenopus* tadpole

**Part I:** The video shows a dorsal view of an isolated *in vitro* preparation of a *Xenopus* tadpole prepared following the protocol in Lambert *et al.* 2008^46^. Ferrofluid was injected into the left inner ear (red-orange color). A permanent point magnet was displaced in proximity above the inner ear. The magnet motion provoked eye rotations via the vestibulo-ocular reflex. Only the left eye responded, as the right eye served to mechanically hold the preparation in place.

**Part II:** (A) Schematic depicting the activation of oblique upward-downward eye movements (curved double arrow) following magnet movement (double arrow) along the ipsilateral posterior vertical semicircular canal (PC) plane of the ferromagnetic particle-filled left inner ear (brown color). (B) Magnet motion-induced a deflection of the PC cupula that in turn causes alternating contractions of the corresponding principal extraocular motor target, the ipsilateral superior oblique eye muscle, thus simulating oblique vestibulo-ocular reflex activity. AC, anterior vertical semicircular canal; HC, horizontal semicircular canal; LR, MR, lateral, medial rectus eye muscle; IO, inferior oblique eye muscle; IR, SR, inferior, superior rectus muscle.Related to Figure 1.

## SUPPLEMENTAL INFORMATION

### Magnetic actuation of otoliths allows behavioral and brain-wide neuronal exploration of vestibulo-motor processing in larval zebrafish

Natalia Beiza-Canelo, Hippolyte Moulle, Thomas Pujol, Thomas Panier, Geoffrey Migault, Guillaume Le Goc, Pierre Tapie, Nicolas Desprat, Hans Straka, Georges Debrégeas, Volker Bormuth

**Figure S1:**
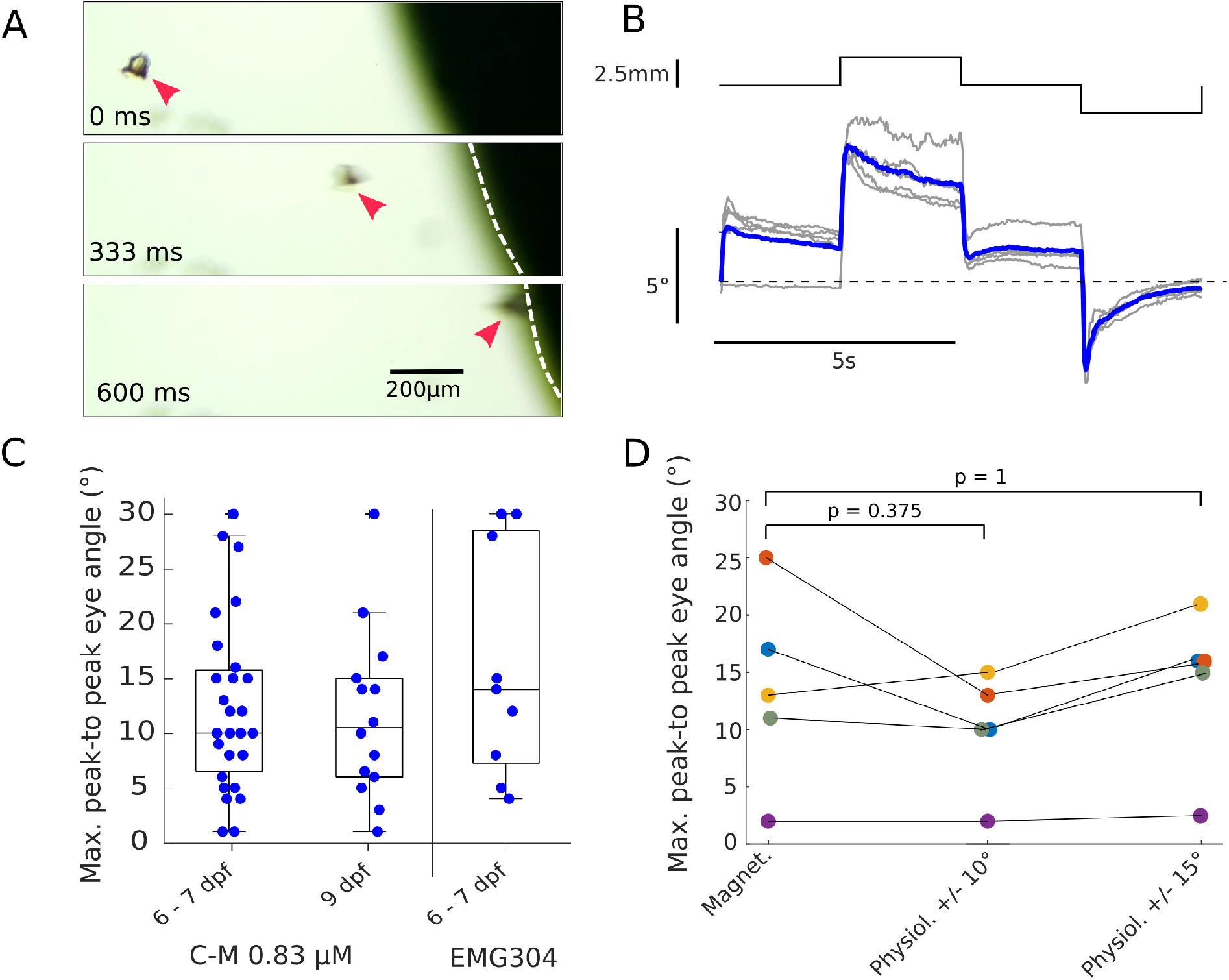
Magnetic actuation of the otoliths after surface coating by ferromagnetic nanoparticles. **(A)** Imaging sequence showing that a ferrofluid coated utricle (red arrowheads), dissected out of the ear, is attracted by a magnet (black, magnet diameter 5 mm, dashed line indicates the magnet’s edge). **(B)** Eye response to a stepwise magnet movement of *±*2.5 mm in amplitude. The response to five consecutive trials is shown in grey and the trial-averaged response is shown in blue.**(C)** Eye rotations evoked in the roll direction by the magnetic stimulation method using the custom ferrofluid injected at 5 dpf and probed at two different time points post injection, 6-7 dpf (N=27) and 9 dpf (N=14). The right boxplot shows the response evoked when using the commercial ferrofluid (see STAR Methods) also injected at 5 dpf and probed at 6-7 dpf (N=9). The means of the three groups were not significantly different (p=0.3973, one-way Anova test). **(D)** Evoked eye rotation angles by the magnetic stimulation methods compared to evoked eye rotation angles evoked by a physiological roll stimulus of +/-10 and +/-15 degrees in amplitude measured sucessively in the same fish (N=5 fish). The Wilcoxon signed rank test was used for statistical comparison. Related to Figure 1.

**Figure S2:**
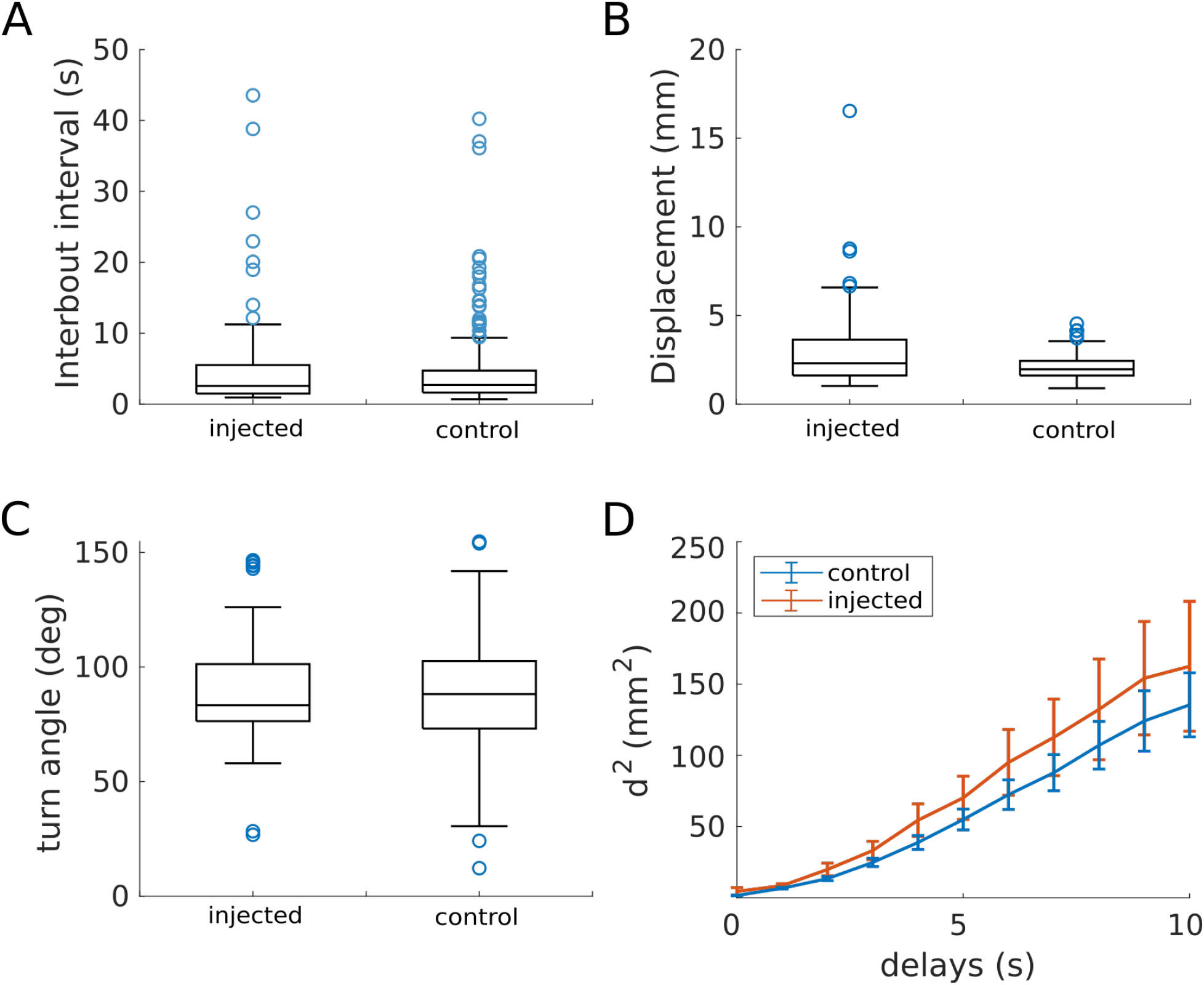
Effect of ferrofluid injection on swimming behavior. **(A-C)** Boxplots showing: **(A)** the distribution of the inter-bout interval, which is the time in seconds elapsed between two consecutive swimming events, **(B)** the mean displacement after a swim bout in mm, and **(C)** the turn angle in degrees. **(D)** Plot showing the mean of the mean square distance for 10 different time delay. The results were obtained by tracking two different batches of injected and non-injected fish. The fish were filmed swimming freely during 1 h (75 fps). P-values: ib-interval p = 0.846, displacement p = 0.00077, turn angle p = 0.366. N = 6 injected fish, 11 control non-injected fish. Related to Figure 2.

**Figure S3:**
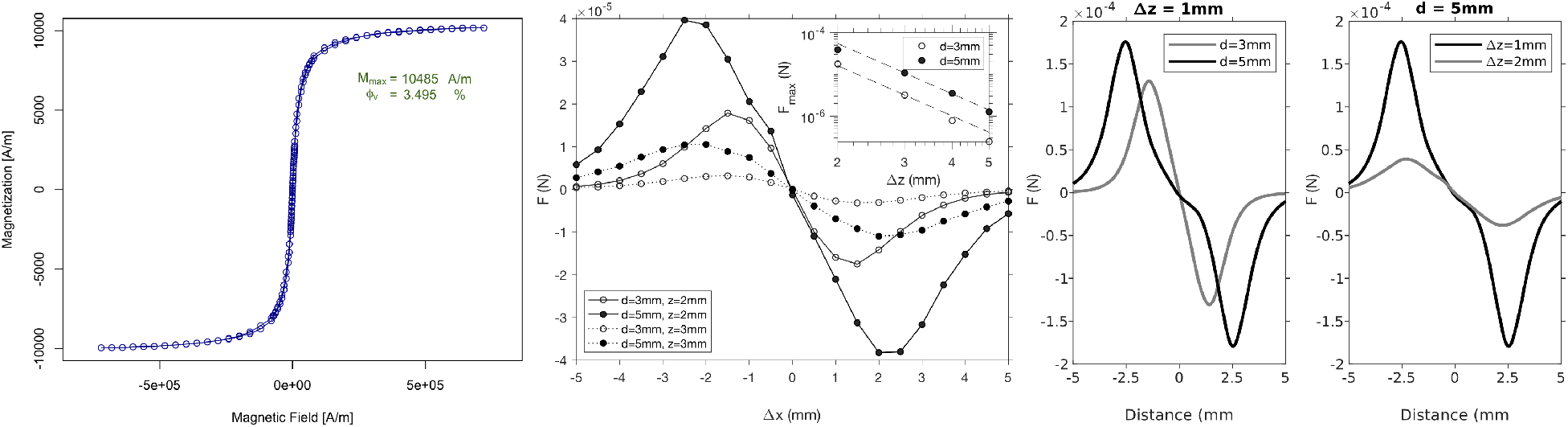
Finite element simulation of the lateral force, F*_L_*, exerted on a paramagnetic particle by a magnet as a function of the lateral distance (Δx) of the particle to the magnet’s center axis. **(A)** Magnetization of the ferrofluid as a function of the magnetic field intensity (H). In vacuum, magnetic field intensity and magnetic flux density are linked by 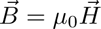. **(B)** Non-normalized simulation results. **(C)** The force-displacement relation becomes more complex when becomes very small compared to the magnet diameter (simulation results are smooths for better visualization). See also figure 13 in Furlani 2012^56^. Related to Figure 3.

